# Digital telomere measurement by long-read sequencing distinguishes healthy aging from disease

**DOI:** 10.1101/2023.11.29.569263

**Authors:** Santiago E. Sanchez, Jessica Gu, Anudeep Golla, Annika Martin, William Shomali, Dirk Hockemeyer, Sharon A. Savage, Steven E. Artandi

**Author notes:** Contributing authors.

## Abstract

Telomere length is an important biomarker of organismal aging and cellular replicative potential, but existing measurement methods are limited in resolution and accuracy. Here, we deploy digital telomere measurement by nanopore sequencing to understand how distributions of human telomere length change with age and disease. We measure telomere attrition and *de novo* elongation with unprecedented resolution in genetically defined populations of human cells, in blood cells from healthy donors and in blood cells from patients with genetic defects in telomere maintenance. We find that human aging is accompanied by a progressive loss of long telomeres and an accumulation of shorter telomeres. In patients with defects in telomere maintenance, the accumulation of short telomeres is more pronounced and correlates with phenotypic severity. We apply machine learning to train a binary classification model that distinguishes healthy individuals from those with telomere biology disorders. This sequencing and bioinformatic pipeline will advance our understanding of telomere maintenance mechanisms and the use of telomere length as a clinical biomarker of aging and disease.

---

Telomeres are nucleoprotein structures at the ends of linear chromosomes which act to prevent the recognition of chromosome ends as double strand breaks and serve to recruit telomerase, the ribonucleoprotein holoenzyme responsible for maintaining telomere length in cells (*1–4*). Telomeric repeats––stretches of non-coding, TTAGGG repeats––are recognized by shelterin, the protein complex (TIN2, TPP1, POT1, TRF1, TRF2, RAP1) that protects telomeres and controls recruitment of telomerase to chromosome ends. In non-malignant cells lacking telomerase, telomeres shorten by several dozens of base pairs during each cell division due to the inability of DNA polymerase to fully replicate the lagging DNA strand (*5,6*). After an extended period of telomere shortening, a subset of the shortest telomeres become dysfunctional or uncapped, signaling a DNA damage response that triggers replicative senescence, autophagy, or apoptosis (*7, 8*). In cells with disrupted Rb and TP53, dysfunctional telomeres precipitate telomere crisis––a process characterized by rampant chromosomal instability stemming from end-to-end fusions at chromosomal termini lacking functional telomeres (*9,10*). The end-to-end fusions and chromosomal instability seen in crisis can be replicated by sabotaging the shelterin complex via deletion or destabilization of TRF2 (*11*). In self-renewing cells––such as stem, germ, and cancer cells––telomerase offsets the end-replication problem by directly elongating telomeres. Telomere maintenance by telomerase depends on recruitment to telomeres by TPP1 and subsequent retention at the single-stranded 3’ overhang end by POT1 (*12*). In cells with sufficiently high telomerase activity, such as embryonic stem cells and cancer cells, this process results in stable telomere lengths over time. In a minority of cancers, an alternative, homologous recombination-dependent telomere lengthening mechanism (ALT) is responsible for maintaining telomere length (*13,14*). Telomeres shorten during physiological aging in most somatic tissues likely due to insufficient telomerase in the stem cell pool (*15–18*). Individuals harboring genetic defects in genes regulating telomere maintenance develop telomere biology disorders (TBD), a group of diseases characterized by aberrantly short telomeres and severe tissue defects (*19–23*). Telomerase-deficient mice display an array of markedly accelerated aging associated phenotypes and have served as models of organismal aging and aging-related disease, including cancer (*8*, *24*). Therefore, telomere length is an important biomarker for the replicative potential as well as the replicative history of a cell in cells with insufficient telomerase to indefinitely maintain telomere length, and, consequently, ensuring telomere maintenance by upregulation of telomerase or initiating ALT is required for any cell aspiring towards indefinite self-renewal.

## Digital telomere measurement by long-read sequencing measures intact telomeres at high-resolution

Telomere lengths have been estimated by Southern blot (TRF) and fluorescence in situ hybridization (FISH) which produce coarse mean telomere lengths by measuring signals from telomeric probes hybridized to genomic DNA. Telomeric probe hybridization has also been leveraged in assays for the measurement and quantification of the shortest telomeres following PCR amplification of chromosome ends (STELA) or adapter-ligated telomere restriction fragments (TeSLA) (*22,23,25*). Recently, it has been demonstrated that long-read sequencing technologies such as PacBio HiFi and Oxford Nanopore (ONT) sequencing can be used to sequence and measure telomeres at unprecedented resolution (*26,27*). We developed a sequencing preparation and bioinformatic pipeline (Telometer) capable of reproducibly measuring telomeres from either whole-genome or telomere-enriched long reads (Fig. 1A). Telomere-containing reads were identified by aligning telomeric repeats to the chromosomal termini of the recently completed telomere-to-telomere human genome (*28*). We defined the extent of each telomere as spanning the distance from the terminal repeat at the chromosome terminus to the final two consecutive repeats preceding the subtelomeric sequence anchoring the telomeric region to its reference chromosome (*26*). The ability to measure the distribution of individual telomere lengths from whole-genome long-read sequencing data enables telomere length analysis to be generically accessible without any additional biochemical intervention prior to sequencing library preparation, but the retrieval of telomeric reads from whole genome sequencing data is highly inefficient due to the paucity of telomere content relative to the rest of the human genome. Capturing the telomeric end with an oligo designed to complement both the telomeric 3’ overhang on one end and the ONT sequencing adapter in combination with restriction digestion of genomic DNA, biochemically enriched for telomeric reads by several thousand-fold without significantly impacting the measurement of the telomere length distribution (Fig. 1B to C). Digital mean telomere length measured by long-read sequencing was highly correlated to existing gold-standards, TRF Southern blot and flow-FISH (Fig. 1D to G). Bootstrapping analysis of our measurement results suggests the standard error of measurement by our method decays exponentially with additional telomere measurements eventually resulting in a maximal precision of 30-40 base pairs (Fig. 1H).

**Figure 1.**
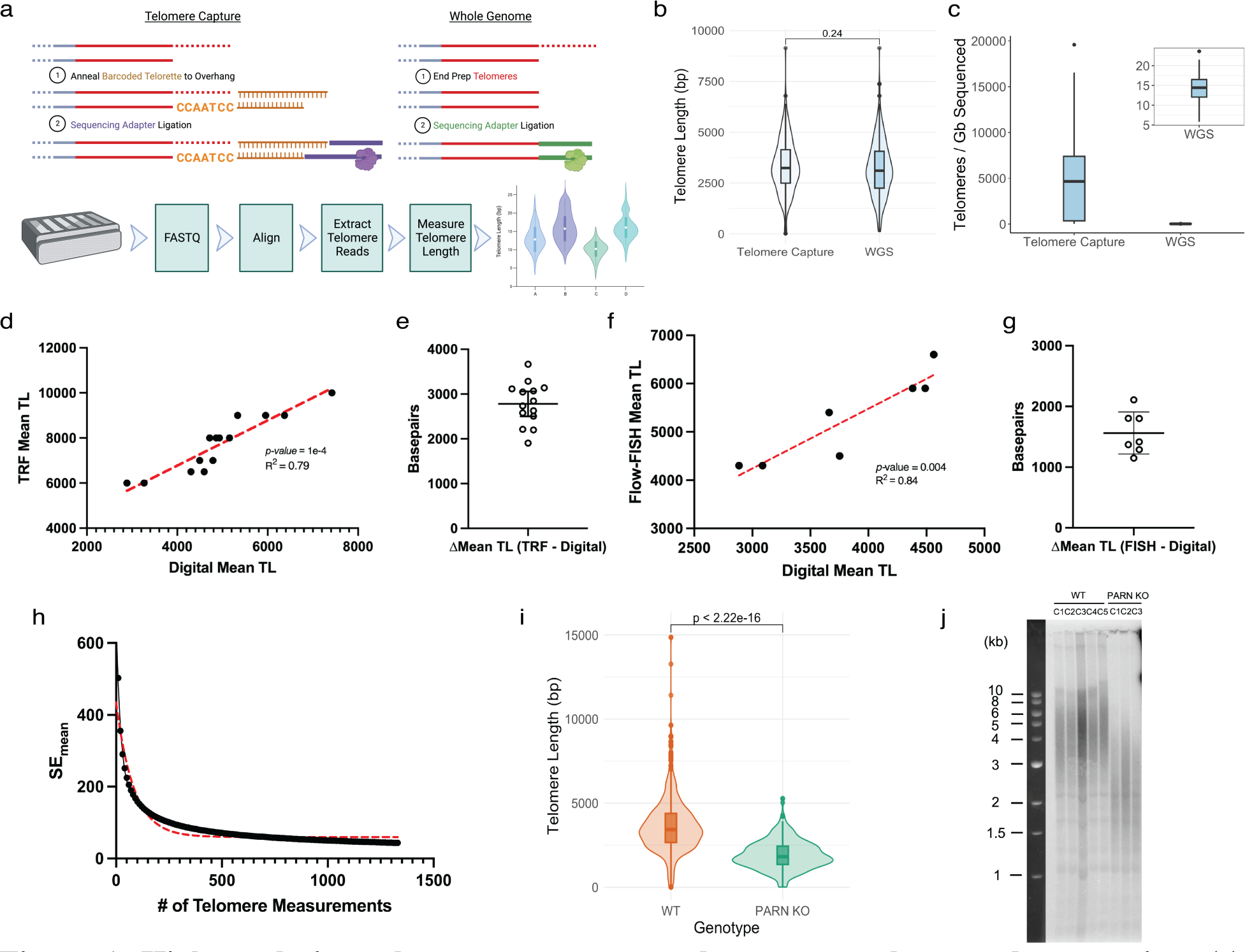
**High-resolution telomere measurement by nanopore long-read sequencing**. (a) Schematic representation of DNA sequencing library preparation for telomere measurement by telomere capture or whole genome long-read sequencing. (b) Head-to-head comparison of telomere length distributions obtained through either telomere capture or whole genome sequencing library preparation from a single source of HEK 293T DNA. (c) Telomeres per gigabase sequenced for both library preparation methods. (d) Correlation between mean telomere lengths from matched samples determined by sequencing or TRF (n=14). (e) Difference in bp between mean telomere length of matched samples measured by both TRF and sequencing (n=14). (f) Correlation between mean telomere lengths from matched samples of RTEL1 mutant individuals determined by sequencing or flow-FISH (n=7). (g) Difference in bp between mean telomere length of matched samples measured by both flow-FISH and sequencing (n=7). (h) Bootstrapping analysis of the change in standard error of the mean telomere length as a function of the total number of telomeres measured. (i) Digital telomere measurement by sequencing of wild-type and PARN KO hESCs (p < 2e-16, Wilcoxon rank-sum test). (j) Analog telomere measurement by TRF of wild-type and PARN KO hESC clones.

To demonstrate whether digital telomere measurement by long-read sequencing can accurately recapitulate known telomere phenotypes, we sequenced several genetically modified human embryonic stem cell (hESC) models of telomere dysfunction including disease-related mutations in PARN or TIN2, and TERT deletion. Genetic defects in PARN––a critical ribonuclease for the appropriate maturation of the human telomerase RNA component (hTR)–– and a gain-of-function mutation (T284R) in TIN2––a shelterin component and negative regulator of telomere length––are two known genetic etiologies of dyskeratosis congenita (DC), a disease characterized by pathologically short mean telomere length in peripheral blood leukocytes (PBLs) (*29–31*). To determine if digital telomere measurement similarly reflects this phenotype, we measured the telomere length distribution of PARN knock-out (Fig. 1I to J), TIN2 T284R heterozygous or homozygous mutants (Fig. 2A), and their corresponding wild-type parental hESCs and found digital telomere measurement can confidently discriminate between wild-type and pathologically short telomere length distributions in hESCs. To understand whether we can measure telomere shortening in human cells lacking telomerase, we sequenced a TERT knockout hESC line (CDKN2A^-/-^ TERT^-/-^, AAVSI:hTERT^flox^), passaged for 66, 78, 98, and 105 days post Cre recombinase-mediated telomerase inactivation and observed progressive shortening between passages (an average of 40 bp per day) (Fig. 2B to C). The shorter mean telomere lengths in genetically edited versus wild-type hESCs is also in broad agreement with analog measurements by Southern blot in these same cells (TRF)(Fig. 1J)(*31*). Finally, to verify that our method captures intact, full-length telomeres we set out to use digital telomere measurement to observe *de novo* telomere addition by forcing a cell line with stable telomeres to overexpress the catalytic core of telomerase (*32*). To this end, we transiently transfected HEK293T cells with a plasmid expressing an hTR encoding a variant telomeric template sequence (TSQ1) or wild-type hTR in addition to TERT, or with GFP alone. We harvested genomic DNA after three days, and performed telomere capture sequencing using capture oligos designed against either the TSQ1 variant sequence (5’-TTGCGG-3’) or the canonical human telomere sequence (5’- TTAGGG-3’). Attempting telomere capture with variant-targeting oligos from genomic DNA harvested from cells not expressing the TSQ1 variant sequence did not produce a successful sequencing library, consistent with an absence of variant telomere repeats in these samples (Fig. S1). Transient transfection with either hTR or TSQ1 and TERT substantially increased *in vitro* telomerase activity and resulted in a 1000 bp increase in the mean telomere length after 3 days in culture (Fig. 2, E to H). Notably, chromosomes with shorter telomeres in the control group (GFP) showed a greater magnitude of elongation relative to chromosomes with longer telomeres when treated with telomerase overexpression (Fig. 2H). The detection of *de novo* telomere elongation by both wild-type and variant capture indicates that our method measures full length telomeres.

**Figure 2.**
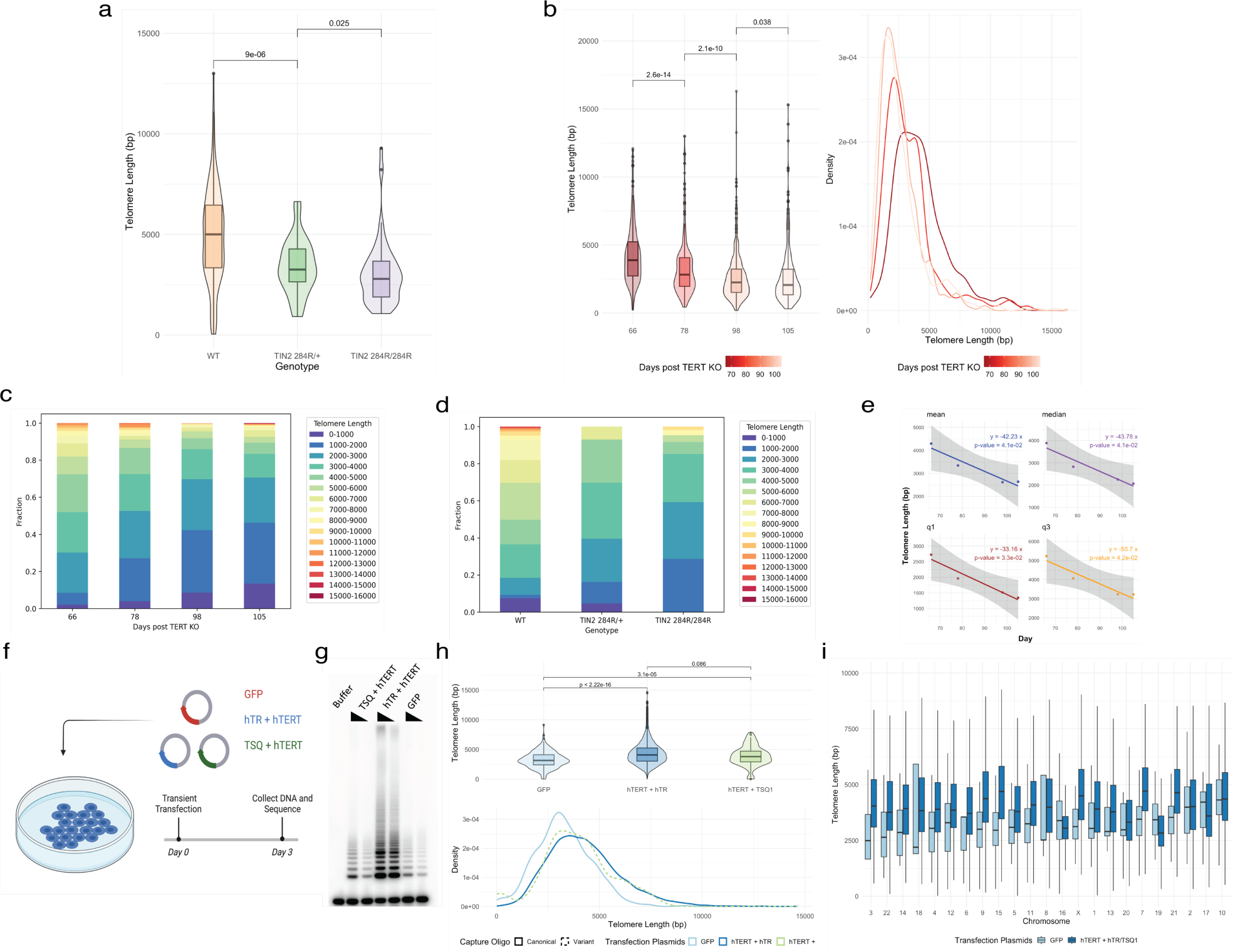
Telomere attrition and *de novo* elongation in cultured human cells. (a) Telomere length distributions of wild-type, 284R heterozygous or homozygous TIN2 mutants. (b) Telomere length distributions of hESCs 66 to 108 days post Cre-mediated TERT knock-out. Box and violin plots of telomere lengths (left) alongside density distributions of telomere lengths (right). (c) Stacked bar graphs of telomere length fractions from hESCs with TIN2 mutations by genotype or (d) days post TERT-knockout. (e) Linear regression of telomere length distribution summary statistics versus days post TERT knock-out. (f) Schematic representation of experiment to measure *de novo* telomere elongation in HEK293T cells. (g) TRAP assay for telomerase activity in transiently transfected HEK293T cells. (h) Telomere length distributions as box or violin plots (Top) and density distributions (Bottom) for transiently transfected HEK293T cells as measured by both canonical and variant telomere capture sequencing. (i) Chromosome-specific telomere length distributions following transient transfection with both hTERT and hTR/TSQ1 or GFP arranged by ascending chromosomal 25^th^ percentile telomere length. In all cases, p-values calculated by Wilcoxon rank-sum test

## High-resolution telomere measurement distinguishes human aging and disease

Mean telomere shortening with age has been observed by previous methods of telomere measurement, however these methods only produce coarse estimates of the population mean (TRF), relative measures of total telomeric content (flow-FISH, qPCR), or rely on PCR to preferentially amplify the shortest telomeres (STELA, TeSLA) (*15–17,21,23*). Moreover, given the resolution of existing methods and the innate variability of telomere length between individuals of similar age, it remains unclear if telomere length can serve as a predictive biomarker of aging. We sequenced DNA derived from PBLs of fourteen healthy human donors aged 18 to 77 years and found that donor age correlated with the mean, median, first quartile, and third quartile telomere lengths Fig. 3A, 3D to E). We observe that the mean and median telomere lengths in peripheral leukocytes decrease by approximately 24 base pairs per year, in close agreement with earlier cross-sectional estimations made from over 1100 TRFs of human PBL DNA (*16*). Strikingly, the third quartile telomere length decreases more steeply with age than the first quartile (Fig. 3E), suggesting that longer telomeres are lost more rapidly than shorter telomeres and echoing observations made in TRF measurements of BJ fibroblasts with limiting telomerase activity in serial passage (*33*). As in BJ fibroblasts, this observation could be explained by insufficient telomerase in the hematopoietic stem cell pool to indefinitely sustain telomere length, or negative selection of cells with increasing fractions of very short telomeres. Quantification of the fraction of telomeres of various sizes in each aging cohort shows that the fraction of the distribution comprised of shorter telomeres increases with age as the longer telomere fraction shrinks with the notable exception of the two shortest fractions which appear to be relatively stable with age in PBLs (Fig. 3H). Our observations quantitatively reinforce a recent observation through TeSLA that telomeres significantly shorter than the mean telomere length accumulate and increase in proportion with age (*22*).

**Figure 3.**
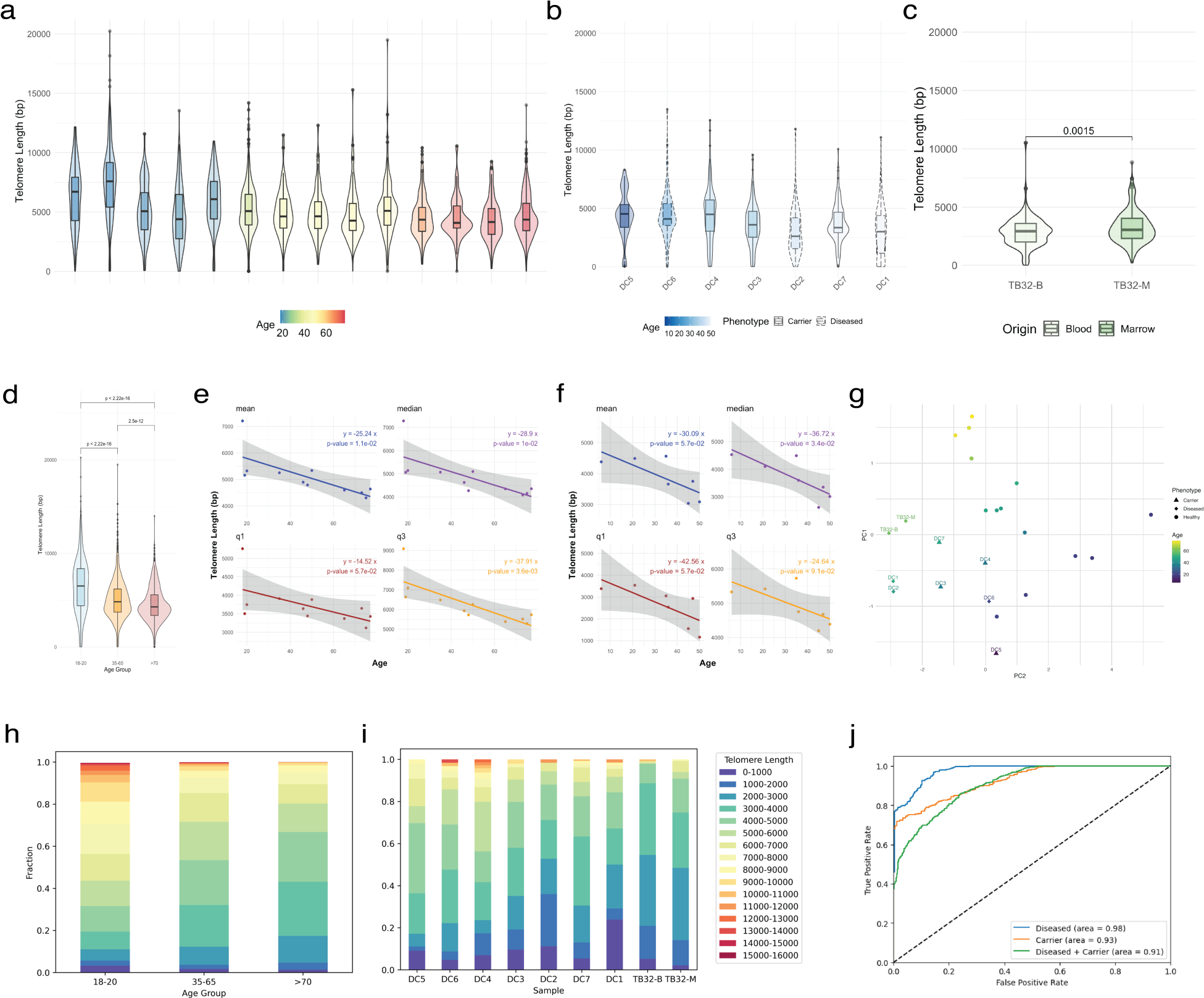
Telomere length distributions distinguish healthy human aging and disease. (a) Telomere length distributions from PBLs of 14 healthy individuals aged 18 to 77 years. (b) Telomere length distributions from PBLs of 7 RTEL1 mutant individuals (healthy carriers = solid outline; diseased carriers = dashed outline). (c) Telomere length distributions from PBLs or DNA obtained from bone marrow biopsy sample of individual under evaluation for a potential telomere biology disorder at Stanford hospital. (d) Telomere length distributions from PBLs of 14 healthy individuals aggregated into young (18-20; n=5), middle (35-65; n=6), and elder (>70; n=3) aged cohorts (*p-*value calculated by Wilcoxon rank-sum test). (e) Linear regressions of telomere length summary statistics versus donor age in 14 healthy individuals (mean = blue, median = purple, q1/25^th^ percentile = maroon, q3/75^th^ percentile = yellow). (f) Linear regressions of telomere length summary statistics versus donor age in 7 samples from RTEL1 mutant individuals (labeled colors as in e). (g) Principal component analysis of telomere length distributions of healthy aging (n=14, unlabeled) and TBD variant carrier samples (n=9, labeled). (h) Stacked bar graph representing telomere length fractions in 1000 bp bins from 0-1000 to 15000-16000 in young, middle, and elder aged cohorts. (i) Stacked bar graph representing telomere length fractions in 1000 bp bins from 0- 1000 to 15000-16000 in 9 samples from individuals with mutations associated with, or being evaluated for, telomere biology disorders. (j) Receiver operator characteristic curve for binary classification of healthy versus carrier, healthy versus diseased, or healthy versus both diseased or carrier telomere phenotype.

To determine if our method is capable of distinguishing healthy aging from telomere biology disorders, we sequenced 7 samples of peripheral blood leukocytes from 6 individuals aged 6 to 50 either diagnosed with TBD or identified as asymptomatic heterozygous carriers of a TBD- associated RTEL1 variant (*22*). In addition, we analyzed genomic DNA from two paired bone marrow (TB32-M) and peripheral blood (TB32-B) samples from one patient at Stanford Hospital with a history of chronic myelomonocytic leukemia (CMML) and pulmonary fibrosis and a newly discovered and previously unreported frameshift mutation in TINF2 (p.Gln298fs) (Table S1). We confirmed telomeres from these individuals were significantly shorter relative to healthy individuals in their age group (Fig. 3B to C, 3F) and observed a significant correlation (R^2^=0.84) with mean telomere length measurements made by flow-FISH, the clinical standard method for telomere measurement in peripheral blood (Fig. 1H). We additionally performed TRF with DNA from twelve of the fourteen healthy donors and the two samples obtained from the Stanford patient (Fig. S2). TRF was found to systematically overestimate mean telomere length by one to three thousand basepairs in all samples for which both digital and analog telomere measurement were performed, most likely due to undigested subtelomeric sequence biasing the results of the Southern blot (Fig. 1F, 1G, S2, S3). Furthermore, while it was not possible to distinguish the telomere lengths from TB32’s blood and marrow samples by TRF, digital telomere measurement revealed the bone marrow telomere length distribution contained longer telomeres than in the same individual’s PBLs (Fig. 3C, S2). An analysis of the telomere length fractions in our cohort of individuals with defective telomere maintenance revealed significant variation between individuals but altogether much higher proportions of the shortest fractions of telomeres compared to healthy donors (Fig. 3H, 3I). Strikingly, TBD patient telomere length distributions were enriched for the two shortest fractions of telomeres (0-2000 bp) relative to healthy donors, suggesting that leukocytes thought to be near the replicative limit can persist in the peripheral blood and that the shortest telomere fractions in the distribution are particularly sensitive to the underlying state of the cell’s telomere maintenance apparatus. If telomerase function in the hematopoietic stem cell compartment is particularly important for preventing the accumulation of the shortest telomere fractions, differential age-associated telomere attrition in telomeres of varying length should be attenuated or disappear in cells with genetic defects in telomere maintenance. In TERT-null hESCs, the daily rate of telomere shortening was at least double the annual age-associated decrease in telomere length for the corresponding distribution statistic in PBLs (Fig. 2B) and the first quartile telomere length in TERT-null hESCs shortened nearly as quickly as the third. In RTEL1 patients and carriers, we no longer observe a difference in the extent of decreasing telomere length in the first and third telomere length quartiles with age (Fig. 3B). It is notable that the TERT-null hESCs we studied reach their replicative limit at 110-120 days post TERT inactivation and nevertheless we observe the relative expansion of the shortest telomere fractions over time in TERT-null hESCs and in RTEL1 patients relative to healthy individuals in similar proportions (*31*). These data suggest that information about the integrity of the underlying telomere length machinery in a cell population can be inferred from the telomere length distribution and its change over time. Furthermore, these data support a model of age-associated telomere attrition where limiting telomerase and not negative selection against cells with very short telomeres is the primary driver behind the accumulation of shorter telomeres with age.

Telomere biology disorders are diagnosed by measuring PBL telomere length by flow- FISH, comparing the result to previously measured population statistics for telomere length for the patient’s age, and subsequent detection of a genetic defect by targeted exon sequencing. Symptomatic patients often have observed mean telomere lengths at or below the first percentile for their age. Telomere biology disorders also exhibit genetic anticipation, and the severity of clinical presentation is inversely proportional to telomere length. As a result, asymptomatic individuals carrying the same genetic defect as affected patients may still have telomere lengths within the lower quintile of the population for their age and subsequently develop symptoms later in life, leading to a delayed TBD diagnosis (*20*,*22*). Digital telomere measurement could also be used, therefore, as a tool in the diagnosis of telomere biology disorders. As a proof of concept, we set out to use machine learning to determine if we could distinguish the following groups by telomere lengths and age alone: symptomatic patients versus healthy donors; unaffected carriers versus healthy donors; or both symptomatic patients and unaffected carriers versus healthy donors. Telomere measurements and ages from our healthy donors, symptomatic patients, or unaffected carriers were randomly and iteratively assigned to serve as training or test data for the binary classification model with a training-to-test ratio of 7 to 3, respectively. In this small sample, our model successfully distinguished both symptomatic and asymptomatic carriers from healthy donors with high sensitivity (Fig. 3J). In the instance of healthy donors versus symptomatic patients, our model achieved 95% accuracy (AUC=0.98); when comparing healthy donors versus asymptomatic carriers, 91% accuracy (AUC=0.93); when comparing healthy donors versus both symptomatic patients and asymptomatic carriers, 86% accuracy (AUC=0.91). Asymptomatic carriers of TBD variants are often not distinguishable from healthy individuals by flow-FISH, but our method could potentially be applied to predict TBD variant carrier status from measurement of the telomere length distribution.

Finally, to validate our observations about the relationship between age, the telomere length distribution, and telomerase activity, we analyzed previously published, high-coverage (30X, on average) whole genome long-read sequencing data from patient-matched colorectal carcinoma and surrounding benign epithelia obtained from twenty individuals predominantly aged between 50 and 70 years (*34*). In this independent dataset, we also observed trends between the four summary statistics of the telomere length distribution and age in benign colonic epithelium, including more rapid shortening of the third quartile relative to the first, but found no correlation between age and any summary statistic of the tumor telomere length distributions, as expected for cancer cells immortalized by telomerase (Fig. 4A to C). We also find that, in this cohort, tumor telomeres were significantly shorter than those measured in benign epithelia in 75% of samples, in agreement with previous studies utilizing TRF, Q-FISH or TelSeq, a short-read sequencing method for estimating relative telomere content, in colorectal carcinoma, melanoma, and TCGA data, respectively (*18,35,36*). Therefore, the telomerase activity present in both hematopoietic and colon epithelial stem cells is insufficient to indefinitely maintain telomere length with the consequence of monotonic shortening, on average, and more rapid shortening of longer relative to shorter telomeres with age in their cellular progeny.

**Figure 4.**
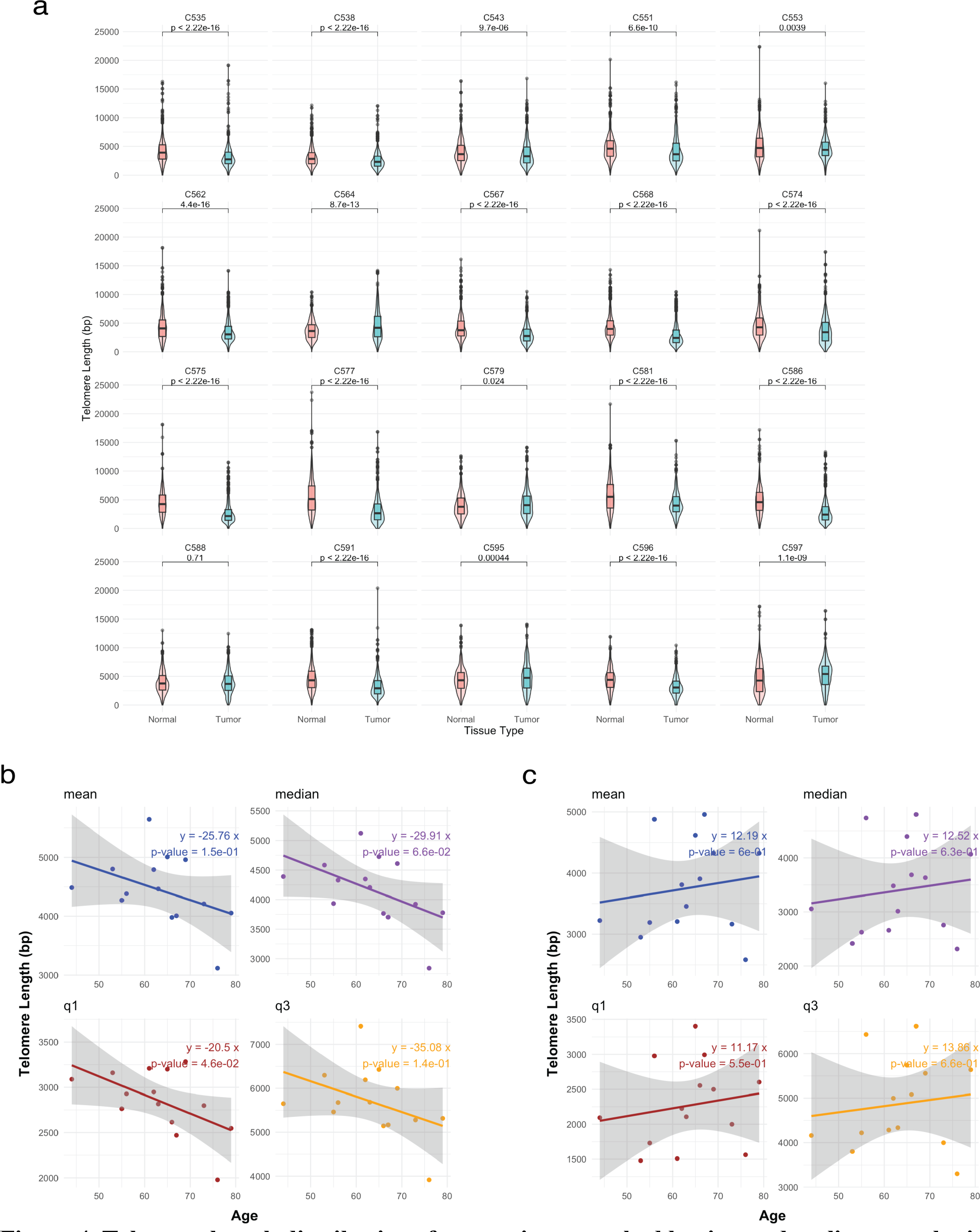
**Telomere length distributions from patient-matched benign and malignant colonic tissue in a cohort of colorectal carcinoma patients**. (a) Grid box and violin plots of telomere length distributions for twenty individuals in the cohort showing matched benign (red) and malignant (blue) colonic tissue by patient ID (*p*-value calculated by Wilcoxon rank-sum test). (b) Linear regression of telomere length distribution summary statistics for benign colonic epithelia versus patient age. (c) Linear regression of telomere length distribution summary statistics for malignant colonic epithelia versus patient age.

## Discussion and Conclusions

For decades, human telomere measurement has only been possible with methods which provided either relative quantifications of telomere content or an estimation of the mean telomere length in a population. Long-read sequencing is revolutionizing our ability to explore the genomic landscape of the chromosomal terminus and we have developed a method which produces high-resolution, high-throughput measurements of individual intact telomeres using nanopore sequencing. In this work, our method recapitulates the dramatic telomere shortening observed in three TBD genotypes both in culture and *in vivo*, *de novo* telomere addition by overexpression of the catalytic core in a cell line with stable telomeres, progressive telomere attrition in human aging and in the presence and absence of functional telomerase. We additionally report a previously undiscovered frameshift mutation in the DC-patch of TINF2 (p.Gln298fs) in a patient with CMML, pulmonary fibrosis, and short telomeres for their age group. Telomere measurements made by long-read sequencing are comparatively much richer than TRF, flow-FISH while requiring less input DNA than either existing technique and are not limited by polymerase processivity like PCR-based methods STELA and TeSLA. Our data show that TRF systematically overestimates mean telomere length by up to several thousand base pairs likely due to the extent of genomic digestion, the restriction enzymes selected for then task, and preparation of the DNA for Southern blot. Flow-FISH better estimates the mean telomere length when compared to TRF, but still overestimates by 1500 bp on average, and flow-FISH’s utility is restricted largely to mean telomere estimation in PBLs. Additionally, the ability to anchor individual telomeres to specific chromosomes enabled us to demonstrate more significant elongation of chromosomes with shorter first quartile telomere lengths prior to overexpression of the telomerase catalytic core, providing novel evidence for the preferential action of telomerase at the shortest telomeres in humans (*8,33,37*). We demonstrate that it is possible to enrich a nanopore sequencing library for telomere-containing reads by several hundred-fold using a combination of custom oligonucleotides to capture telomeric ends and restriction digestion to reduce non-telomeric DNA. Nevertheless, our bioinformatic pipeline, Telometer, can be generically applied to any set of human whole-genome long-read sequencing data produced by either ONT or PacBio sequencers.

Many studies, including this one, have demonstrated that the mean human telomere length decreases with age. In this study, we examine telomere length distributions from a cross-section of 63 healthy and diseased human samples and demonstrate the potential of digital telomere measurement as a tool for clinical investigation. We make the additional novel observation that the structure of the telomere length distribution––interquartile telomere lengths, median, mean, fraction of telomeres of varying length––contains information about the underlying telomere maintenance mechanism of the cell. We also find that the association between age and length in telomeres of different length can vary substantially in some tissues, for example between the first and third quartiles of the length distribution. Our data suggest that the shorter end of the telomere length distribution is sensitive to the function of telomerase, and inactivating telomerase *in vitro* or impairing its function *in vivo* both have a marked impact on the maintenance of the lower end of the telomere length distribution. In the two compartments we study (peripheral blood and colonic epithelia) in aging cohorts, there appears to be some variation even in healthy aging-associated telomere attrition between the two compartments. This difference could be explained by the dosage of telomerase, the proliferation rate, or perhaps even the dosage of shelterin components within the stem cell compartments of each tissue, but more comprehensive investigation of longitudinal telomere length evolution in healthy aging across various tissues is required. Generating ensembles of high-resolution measurements instead of means from a single noisy signal permitted us to train a binary classification model by machine learning capable of distinguishing healthy and diseased telomere distributions. Notably, although asymptomatic carriers are not always readily distinguishable from normal individuals by flow-FISH, we demonstrate our simple binary classification model is capable of doing so with high sensitivity. Long-read sequencing technologies are already being deployed in the clinical setting for the rapid diagnosis of genetic disorders and as a tool in clinical research (*38*), and this work demonstrates that digital telomere measurement holds promise as a diagnostic or prognostic tool for the evaluation of telomere biology disorders. As the repertoire of long-read genomic data grows, analysis of telomere length distributions from larger, more diverse, and ideally longitudinal studies will enable telomere length to become a more robust and perhaps even predictive biomarker of aging and disease.

Surprisingly, in every sequencing experiment we have performed in either humans or cultured cell lines, including embryonic stem cells, we have measured telomeres as short as dozens of base pairs. Our work demonstrates that the shortest telomere fractions are sensitive to functional perturbation, suggesting that even if some extremely short telomere measurements are somehow artefactual, cells can tolerate some quantity of very short telomeres without triggering replicative senescence or experiencing chromosomal instability, likely due to the critical protective role of shelterin at telomeres. Future work leveraging digital telomere measurement could elucidate the critical mass of very short telomeres or the thresholds at which either shelterin must be impaired or telomeres sufficiently shortened prior to triggering replicative senescence or observing telomere fusions. Access to the complete telomere length distribution of a cell also opens the possibility of more effectively studying the effects of perturbing the cell’s telomeric rheostat (*39*) on telomere length. For example, in this work we find that in a previously studied cohort of patient-matched colorectal carcinoma and benign colonic epithelia, tumor cells harbored shorter telomeres in approximately 75% of patients. Since telomere length distributions in tumors are known to be stable over time, it follows that the setpoint of a tumor’s telomeric rheostat is somehow related to its unique cellular biology but until now it has not been possible to assess precisely how a certain rate of division, telomerase activity, and concentration of key telomere maintenance machinery like the RNA template, telomerase holoenzyme, or shelterin component proteins individually or quantitatively contribute to stable telomere length distributions. Understanding this dynamic biochemical equilibrium could reveal novel regulators of telomere length as well as new approaches towards therapeutically targeting the telomere maintenance machinery in cancer.

## Online Methods

### High-molecular weight DNA isolation and quantification

High-molecular weight (HMW) DNA was extracted using the NEB Monarch HMW DNA Extraction kit for Cells and Blood (Catalog #T3050L) according to the kit manufacturer’s instructions. Briefly, for cells maintained in tissue culture, cells were trypsinized until detached and then centrifuged at 1000 g for three minutes before adding prep and lysis solution according to the manufacturer’s instructions. Cells were incubated for 10 minutes at 1800 RPM in a thermomixer and DNA from the lysed cells was precipitated onto glass beads, washed twice with 80% ethanol, and finally eluted in Monarch Elution Buffer II according to manufacturer’s instructions. DNA was quantified using a Qubit 4 fluorometer and the Qubit BR dsDNA quantification reagents (Catalog #Q32850). For DNA extraction from peripheral blood, red blood cell lysis was first performed prior to DNA extraction according to the manufacturer’s instructions. DNA quality was assessed by Nanodrop and the average molecular weight was verified to be 60 kb or larger using an Agilent Tapestation (Catalog #5067-5365, 5067-5365).

### Telomere Restriction Fragment Southern Blot

Approximately 4 µg of genomic DNA was prepared in a 50 uL total volume restriction digest solution (1X Fast Digest Buffer, 3 µL HinfI, and 3 µL RsaI) and allowed to digest at 37°C overnight. In the morning, 1 µL each of HinfI and RsaI was added to each digestion reaction and allowed to incubate at 37°C for a further three hours. A 1% TAE agarose gel was prepared and 3 µL per sample underwent gel electrophoresis (125V, 70 minutes) to confirm restriction digest completed successfully (Figure SX). A 0.8% TBE agarose gel was then prepared in a 20x27 cm casting tray after adding 15 µL ethidium bromide to the agarose solution. The entire volume of each restriction digest reaction was then loaded into each well with 1X NEB nucleic acid loading dye in addition to NEB 1 kb reference ladder and gel electrophoresis was performed (85V for 16 hours). In the morning, the gel was dried using a BioRad gel dryer (1 hour under vacuum then 1 hour under vacuum and 50°C). A UV-translucent ruler was then overlayed on the dried gel before imaging with UV transillumination to establish reference distances for the ladder markers from the well positions. The dried gel was then incubated in denaturing buffer (1.5M NaCl, 0.5M NaOH) for one hour with gentle shaking. The denatured gel was washed with deionized water twice before a second one-hour incubation in neutralizing buffer (1.5M NaCl, 1M Tris-HCl, pH 7.4) with gentle shaking. The neutralizing buffer was decanted and the neutralized gel was washed twice with deionized water. The gel was then rolled vertically into a glass hybridization tube (Thermo-Fischer Scientific) and incubated with pre-warmed hybridization buffer (Invitrogen #AM8670) at 42°C for 30 minutes with rotation. 0.5 µM of γ-^32^P labeled telomere probe was then added to the hybridization buffer tube and incubated at 42°C overnight with rotation. The gel was washed once with 2X SSC buffer and twice more with 1X SSC buffer (0.15M NaCl, 15 mM sodium citrate) before exposing onto a phosphor screen inside a lead exposure cassette for 24 hours. Following exposure, the phosphor screen was imaged on a Typhoon scanner.

Both the southern blot images and the ethidium bromide reference ladder were loaded onto ImageJ and aligned. The signal intensities at each position coordinate starting from the bottom of the well in the southern blot image were obtained with the ImageJ line and Measure tools after drawing a line from the bottom of the well to the bottom of the gel through each sample lane.

### Telomerase repeated amplification protocol (TRAP)

To measure telomerase activity, a two-step TRAP procedure was performed as previously described (*36*). Briefly, cell protein extracts (at 1X or 3X dilution with lysis buffer per transient transfection condition) were incubated with telomeric primers for 30 min at 30°C in a PCR machine, followed by 5 min of inactivation at 72°C (cold extension). 1 μl of the cold extension reaction was PCR amplified (24 cycle of 30 s at 94°C, followed by 30 s at 59°C) in the presence of ^32^P end-labeled telomeric primers. The radiolabeled PCR reactions were resolved by 9% polyacrylamide gel electrophoresis at room temperature, and the gel was exposed to a phosphor- imager and the phosphor screen was scanned by a Typhoon scanner.

### Whole-genome sequencing nanopore library preparation

Nanopore library preparation for whole-genome sequencing was carried out according to Oxford Nanopore Technologies (ONT) protocol for native genomic DNA sequencing (LSK-110) with some modifications. Briefly, approximately 1 µg of DNA per sample was end-prepped using the FFPE DNA repair and Ultra II End-Prep enzyme mixes from the NEBNext companion module for ONT ligation sequencing (Catalog #E7180L). The end-prep reaction was incubated in a thermocycler at 20°C for 30 minutes and then 65°C for 30 minutes. End-prepped DNA was extracted from the reaction using Promega ProNex size selection beads (Catalog #NG2001) at a bead-to-reaction solution ratio of 1.6 and incubated on a Hula mixer at room temperature for five minutes prior to being pelleted on a magnet and then washed twice with 80% ethanol and then allowed to dry on the magnet for 3 minutes. DNA was eluted from the beads using ONT elution buffer at 37°C for 15 minutes. For simplex experiments, sequencing adapters (ONT AMXF) were then ligated to the end-prepped DNA for one hour at room temperature using NEB Quick T4 DNA ligase (Catalog #E7180L). For multiplex experiments, ONT barcodes were ligated using NEB Blunt/TA ligase for one hour at room temperature and barcoded DNA was extracted, pooled up to 1 µg of total DNA, and then ligated to sequencing adapters as described previously. Adapter-ligated DNA was extracted from the ligation reaction using Promega ProNex size selection beads at a bead-to-reaction solution ratio of 1.1 and incubated and eluted as described previously. 20-50 fmols of Adapter-ligated DNA was sequenced on R9.4.1 PromethION flow cells on a P2Solo for 24-72 hours. It is advisable to quantify DNA after every bead purification step using a Qubit fluorometer and the Qubit dsDNA BR DNA quantification assay.

Genetically modified hESCs were sequenced on R10.4 PromethION flow cells on a P2Solo and therefore the sequencing adapter used was changed to pair with the updated flow cell chemistry, per ONT’s standard sequencing protocol (adapter NA instead of AMII for telomere capture).

### Telomere capture sequencing nanopore library preparation

Barcoded telomere capture oligos were annealed to sequencing tether (seqTether) by mixing equimolar amounts of both oligos in low TE buffer, heating to 95°C for 2 minutes and then being allowed to cool at room temperature for an hour. Approximately 3 ug of HMW genomic DNA was ligated to barcoded, freshly duplexed oligos in a 100 µL ligation reaction (10 µL 10X rCutSmart buffer, 5 µL 5 µM duplex capture oligos, 2 µL 2000U/µL T4 DNA ligase, 1 µL 10 mM ATP, 3 µg gDNA, nuclease-free H2O up to 100 µL) overnight at 37°C. The following day the ligation reaction was heat inactivated at 65°C for 10 minutes. Potential gaps between the capture oligo and the double-strand/single-strand junction were then filled in using the same reaction tube by adding 2 µL (4U) *Sulfolubus* DNA Polymerase IV (NEB #M0327S), 12 µL 10X ThermoPol Buffer, 1 µL 20 mM dNTPs, 1 µL 10 mM ATP, and 4 µL of nuclease-free H2O followed by incubation at 56°C for 2 minutes and then 72°C for 15 minutes with shaking (500 RPM). Promega ProNex size selection beads were then added to the ligation reaction at a bead-to-solution ratio of 1.6 and the solution was equilibrated on a rotating mixer for 5 minutes at room temperature. The bead solution was then pelleted on a magnet, washed twice with 80% ethanol, allowed to dry on the magnet for 3 minutes, and then eluted in 60 µL of ONT elution buffer for 15 minutes at 37°C. The complete volume of eluted capture-oligo ligated DNA was then ligated onto the ONT sequencing adapter (AMII for R9, NA for R10 libraries) per ONT’s barcode ligation sequencing protocol (5 µL sequencing adapter mix, 20 µL 5X Quick T4 DNA Ligase Buffer, 10 µL Quick T4 DNA Ligase, 5 µL nuclease-free H2O) at room temperature for one hour. The rest of the library preparation and sequencing protocol is performed as for whole-genome sequencing above.

### Data Analysis

Sequencing data was basecalled using Guppy 6.3.0 (ONT) high-accuracy basecalling and aligned to a human telomere-to-telomere reference genome (T2T-CHM13 + Stong 2014 subtelomere assemblies (*41*)) with minimap2. Aligned BAM files were then sorted and indexed with samtools (v.1.16.0) and telomere measurements were extracted from alignments using a custom script (TBD). Telometer is based on previously published work for telomere measurement from PacBio HiFi reads (*26*) which was adapted for telomere measurement from Oxford nanopore long reads. In brief, reads mapping to the terminal arms of each chromosome are searched for telomeric repeats by using regular expressions targeted at the sequence patterns corresponding to both the canonical human telomeric repeat (TTAGGG/CCCTAA) and the commonly miscalled motifs previously identified in the literature (*42*). A check is then performed to ensure that identified telomeric sequences are terminal and telomeres are measured from the read terminus until the sub/telomeric boundary. Conceptually, the sub/telomeric boundary is encountered by moving from the telomere terminus inward until telomeric repeats give way to non-telomeric sequence. We formally define the sub/telomeric boundary as the genomic position corresponding to the starting position of the final two telomeric repeats before encountering non-telomeric motifs. The telomere length is then the length, in basepairs, spanned by consecutive telomeric sequence motifs at the terminus of chromosome arms. Figures were made and statistical analyses performed with R (v4.1.0).

For samples sequenced on R10.4 PromethION flow cells, raw pod5 files were basecalled using a custom bonito telomere calling model (HG002.k1) provided by Oxford Nanopore Technologies. As the R10 nanopore chemistry differs significantly from R9, the raw signal from the flow cell is substantially changed and telomere identification from basecalled reads with our custom scripts is not possible with data produced from the default guppy R10 basecaller; however, ONT’s custom telomere basecalling model once again makes it possible for our algorithm to accurately identify telomeric reads from R10 data. Training ONT basecalling models to basecall telomeres more accurately has been previously demonstrated in the literature (*42*). Since work for this manuscript was completed, ONT has incorporated improved telomere basecalling into the default dorado basecaller (v0.3.4, dna_r10.4.1_e8.2_400bps_sup@v4.2.0). Telomere capture experiments with all three basecalling models on both sequencing chemistries are compared in Figures S7 and S8. Besides the supplementary basecaller comparison, all experiments in the same figure were performed on the same sequencing chemistry and basecalled with the same model. Otherwise, library preparation and data analysis were performed as described above.

### Binary Classification Model

To develop a predictor of the disease or carrier status of patients based on telomere length and other minimal sequencing data, the Multi-Layer Perceptron Classifier (MLPClassifier) from Scikit-learn 1.2.2 was employed. The MLPClassifier is a feedforward artificial neural network model that maps input data sets to a collection of acceptable outputs. An MLP is made up of numerous layers, each of which is completely connected to the one before it. Except for the input layer, the remaining neurons have nonlinear activation functions. The MLPClassifier used for this study contained three hidden layers of 8 neurons each, the ‘relu’ activation function applied sequentially, the ‘adam’ solver, a maximum of 500 iterations, and early stopping engaged. Two types of classification were pursued: diseased vs. healthy and both diseased and carrier patients vs. healthy donors. The ‘telomere length’, ‘chromosome’, ‘age’ data fields from Telometer output were provided as input to the classifier and the individuals ‘phenotype’ was used as the classification output (Fig. S9). To train both models, a 70:30 training:test data split was used, and randomized validation sets were used during training for validation score calculations at each iteration (Table S3).

### Transient Transfection of HEK293T Cells

HEK293T cells were cultured in DMEM supplemented with 10% FBS and 1% penicillin/streptomycin. Transient transfection was carried using a 3:1 polyethylenimine to plasmid ratio and 3 µg of DNA, equal parts pCDNA-2xStrep-3xFLAG-hTERT and either pBS- U1-hTR or pBS-U1-TSQ1 containing plasmids. Following the addition of PEI and transfection plasmids, cells were maintained in culture for three days. Cells were then trypsinized and collected and approximately one-million cells were set aside for downstream interphase DNA FISH. High molecular weight DNA was collected from the remaining cells for each condition according to the NEB Monarch High Molecular Weight DNA Extraction Kit for Cells and Blood (NEB #T3050L) protocol. Extracted DNA was quantified using the Qubit BR dsDNA assay (Invitrogen #Q32853) and a Qubit 4 fluorometer.

Frozen pellets of human embryonic stem cells for TERT knock-out and TIN2 mutant hESCs were generously donated by Dirk Hockemeyer’s laboratory and immediately processed for sequencing without restarting culture.

## Supplementary Information

This article contains supplementary information.

## Acknowledgments

We are grateful to the participants in the NCI’s Inherited Bone Marrow Failure Syndrome study, without whom this work would not be possible. We thank Dr. Neelam Giri, NCI, for clinical support. We also thank the Stanford Blood Center and the several blood donors whose donations contributed to this work under IRB #13942. We thank Oxford Nanopore Technologies for providing development access to their custom telomere basecalling model for R10 flow cell chemistry sequencing data. We thank the entire Artandi laboratory for their discussions and support. We also thank Tobias Schmidt and Jan Karlseder (Salk Institute) for coordinating manuscripts and sharing their insight.

## Contributions

SES and SEA conceived the project and methodology. SES designed the experiments, wrote Telometer, analyzed the data. JG cultured and provided PARN WT and KO hES cells and performed the TRF Southern blot of PARN WT and KO hES clones. AM cultured, prepared, and provided TIN2 mutant and TERT knock-out hES cells in the laboratory of DH. SAS provided peripheral blood DNA from RTEL1 variant cohort and had previously consented and participated in clinical care for these patients. SAS performed flow-FISH and TeSLA on RTEL1 variant cohort DNA and provided access to subsequent data analysis. WS participated in the clinical care of TB32, consented the patient for peripheral blood and bone marrow sample collection, and provided the flow-FISH and exon sequencing results for TB32. AG trained and tested the binary classification model. SES and SEA wrote the original draft of the manuscript and all authors participated in subsequent review and editing.

## Data availability

Sequencing data from cell lines is available the NCBI Sequence Read Archive at accession number TBD. For privacy reasons, human sequencing data is available by specific request to the NCBI’s database of Phenotypes and Genotypes at TBD. Our bioinformatic pipeline for telomere measurement from nanopore long-reads is available on github at TBD. Pre-trained binary classification models and code used to train them is also available on github at TBD.

