## Supplementary Information for "Digital telomere measurement by long-read sequencing distinguishes healthy aging from disease"

**Extended Data and Figures**

**Supplementary Table 1A. Characteristics of healthy peripheral blood donors, RTEL1 mutants, and TB32**

TBD Variant Cohort

| UPN (*22*) | Sample ID | Gene | Variant | Age | Notes |
|---|---|---|---|---|---|
| NCI-297-4 | DC1 | RTEL1 | c.1338+3 A>G (IVS15+3 A>G) | 50 | Same affected individual, sampled 5 years apart |
| | DC2 | | | 45 | |
| NCI-297-5 | DC7 | RTEL1 | c.3361delG (p.A1121LfsX6) | 47 | healthy carrier |
| NCI-297-3 | DC6 | RTEL1 | c.1338+3 A>G (IVS15+3A>G) and c.3361delG (p.A1121Lfs*6) | 21 | affected biallelic |
| NCI-345-4 | DC3 | RTEL1 | c.3442del (p.His1148Thrfs*12) | 36 | healthy carrier |
| NCI-345-3 | DC4 | RTEL1 | c.49C>T (p.Pro17Ser) | 35 | health carrier |
| NCI-180-2 | DC5 | RTEL1 | c.1845G>T (p.E615D) | 6 | healthy carrier |
| | TB32 | TINF2 | c.891_892insTTGCTCT (p.Gln298fs) | 60 | affected |

Healthy Donors (Stanford Blood Center, IRB#13942)

| Sample ID | Age | Sample ID | Age |
|---|---|---|---|
| ma01 | 46 | oa60 | 77 |
| ma04 | 65 | oa68 | 75 |
| ma06 | 50 | ya40 | 20 |
| ma13 | 46 | ya41 | 18 |
| ma20 | 48 | ya45 | 18 |
| ma66 | 35 | ya55 | 19 |
| oa51 | 73 | ya56 | 20 |

**Supplementary Table 1B. Sample ID and age at biopsy of patient-matched benign colonic epithelia and tumor biopsies.**

CRC Cohort (34) N=benign/normal; T=tumor

| ID | Age | ID | Age | ID | Age | ID | Age |
|---|---|---|---|---|---|---|---|
| C535-N | 73 | C575-N | 55 | C562-N | 63 | C588-N | 66 |
| C535-T | 73 | C575-T | 55 | C562-T | 63 | C588-T | 66 |
| C538-N | 76 | C577-N | 61 | C564-N | 67 | C591-N | 63 |
| C538-T | 76 | C577-T | 61 | C564-T | 67 | C591-T | 63 |
| C543-N | 55 | C579-N | 79 | C567-N | 62 | C595-N | 56 |
| C543-T | 55 | C579-T | 79 | C567-T | 62 | C595-T | 56 |
| C551-N | 69 | C581-N | 62 | C568-N | 55 | C596-N | 44 |
| C551-T | 69 | C581-T | 62 | C568-T | 55 | C596-T | 44 |
| C553-N | 65 | C586-N | 53 | C574-N | 62 | C597-N | 67 |
| C553-T | 65 | C586-T | 53 | C574-T | 62 | C597-T | 67 |

**Supplementary Table 2. Telomere Capture Oligo Sequences**

| | |
|---|---|
| seqTether | AACCTTGGAGATGCACGGAGCAAGCAAT |
| | **Barcoded Canonical Telomere Capture Oligos (Modification: 5' Phosphate)** |
| p-t3-nb01 | TGCTCCGTGCATCTCCAAGGTTCACAAAGACACCGACAACTTTCTTCCTAACC |
| p-t3-nb02 | TGCTCCGTGCATCTCCAAGGTTACAGACGACTACAAACGGAATCGACCTAACC |
| p-t3-nb03 | TGCTCCGTGCATCTCCAAGGTTCCTGGTAACTGGGACACAAGACTCCCTAACC |
| p-t3-nb04 | TGCTCCGTGCATCTCCAAGGTTTAGGGAAACACGATAGAATCCGAACCTAACC |
| p-t3-nb05 | TGCTCCGTGCATCTCCAAGGTTAAGGTTACACAAACCCTGGACAAGCCTAACC |
| p-t3-nb06 | TGCTCCGTGCATCTCCAAGGTTGACTACTTTCTGCCTTTGCGAGAACCTAACC |
| p-t3-nb07 | TGCTCCGTGCATCTCCAAGGTTAAGGATTCATTCCCACGGTAACACCCTAACC |
| p-t3-nb08 | TGCTCCGTGCATCTCCAAGGTTACGTAACTTGGTTTGTTCCCTGAACCTAACC |
| | **NB01-Barcoded Variant Telomere Capture Oligos (Modification: 5' Phosphate)** |
| p-tsq1-nb01 | TGCTCCGTGCATCTCCAAGGTTCACAAAGACACCGACAACTTTCTTCGCAACC |
| p-tsq2-nb01 | TGCTCCGTGCATCTCCAAGGTTCACAAAGACACCGACAACTTTCTTCCGCAAC |
| p-tsq3-nb01 | TGCTCCGTGCATCTCCAAGGTTCACAAAGACACCGACAACTTTCTTCCCGCAA |
| p-tsq4-nb01 | TGCTCCGTGCATCTCCAAGGTTCACAAAGACACCGACAACTTTCTTACCCGCA |
| p-tsq5-nb01 | TGCTCCGTGCATCTCCAAGGTTCACAAAGACACCGACAACTTTCTTAACCCGC |
| p-tsq6-nb01 | TGCTCCGTGCATCTCCAAGGTTCACAAAGACACCGACAACTTTCTTCAACCCG |
| p-tsq7-nb01 | TGCTCCGTGCATCTCCAAGGTTCACAAAGACACCGACAACTTTCTTGCAACCC |

**Supplementary Table 3. Binary classification model training and test results**

| | Diseased | Carrier | Diseased + Carrier |
|---|---|---|---|
| Training Accuracy | 0.96 | 0.92 | 0.86 |
| Test Accuracy | 0.95 | 0.91 | 0.86 |

| | 1 | 0 |
|---|---|---|
| Model 1 | Diseased | Not Diseased |
| Model 2 | Carrier | Not Carrier |
| Model 3 | Diseased or Carrier | Neither Diseased nor Carrier |

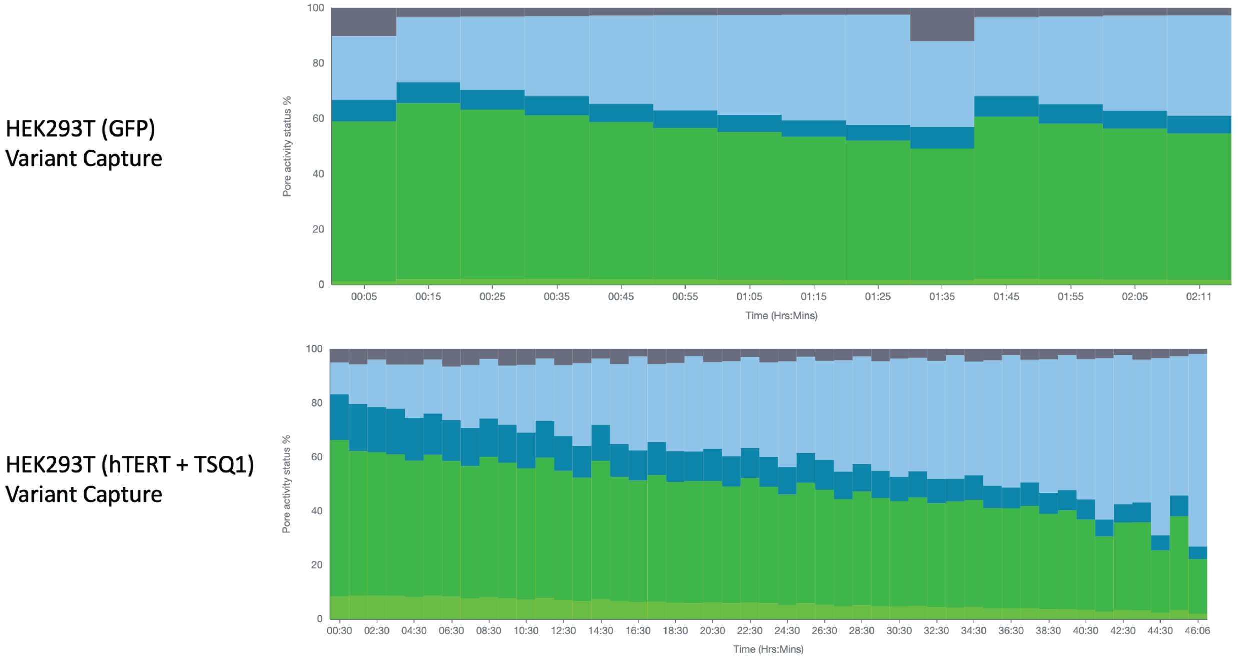


**Fig. S1. Variant telomere capture only produces successful libraries in HEK293T cells transiently transfected with a variant telomerase RNA component template (TSQ1).** Pore occupancy vs. time as produced by ONT MinKNOW software during sequencing. Dark green bars represent pores available for sequencing, light green represent actively sequencing pores.

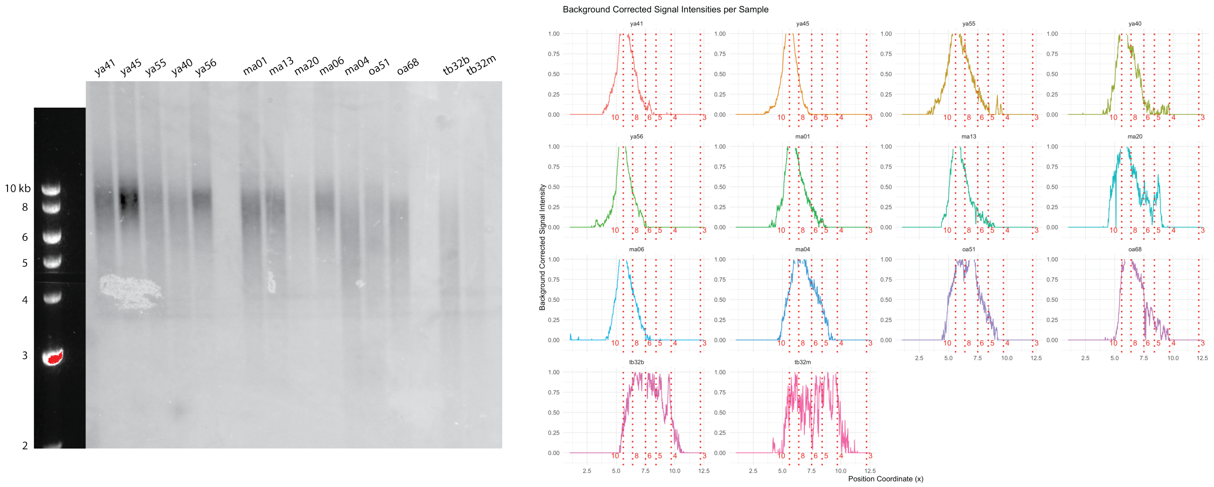


**Fig. S2. Telomere restriction fragment Southern Blot analysis of twelve healthy donors and Stanford Hospital patient.** (left) Phosphor image of TRF from twelve healthy donors and Stanford hospital patient aligned to ethidium bromide image of corresponding DNA ladder from gel. (right) Quantitative analysis of signal from TRF Southern Blot using R (4.1.0), analysis code available on github.

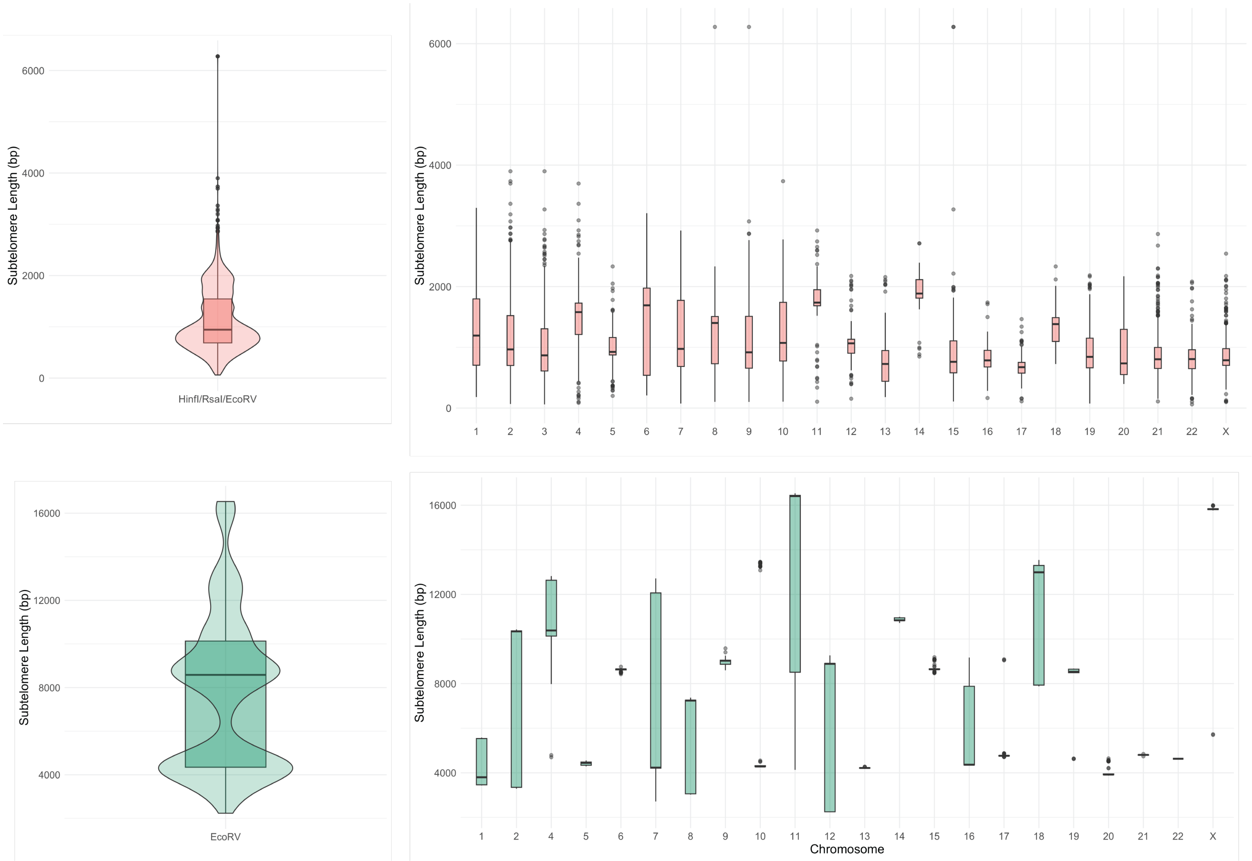


**Fig. S3. Subtelomere lengths measured from telomeric reads in samples digested by single or combination restriction enzyme digest.** (top) Subtelomeric lengths, aggregate and chromosome-specific (left, right, respectively) measured in telomeric reads from previously published PacBio data sequencing DNA previously digested by HinfI/RsaI/EcoRV (*12*). (bottom) Subtelomeric lengths, aggregate and chromosome-specific (left, right, respectively) measured in telomeric reads from nanopore sequencing in DNA previously digested with EcoRV alone (original data).

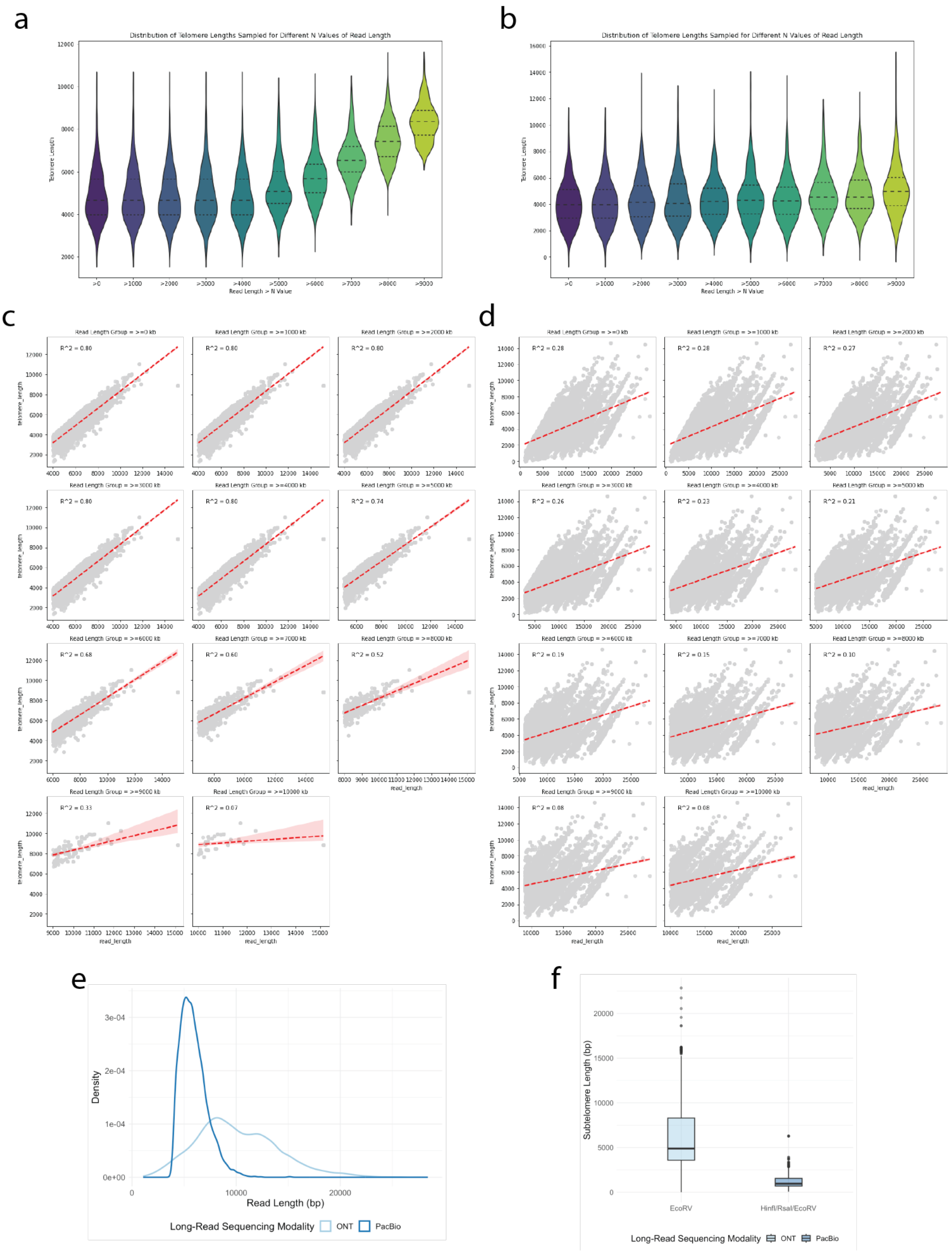
a
Distribution of Telomere Lengths Sampled for Different N Values of Read Length
Telomere_Length
Read Length > N Value
>0
>1000
>2000
>3000
>4000
>5000
>6000
>7000
>8000
>9000
b
Distribution of Telomere Lengths Sampled for Different N Values of Read Length
Telomere_Length
Read Length > N Value
c
Read Length Group = >=0 kb
R^2 = 0.80
Read Length Group = >=1000 kb
R^2 = 0.80
Read Length Group = >=2000 kb
R^2 = 0.80
Read Length Group = >=3000 kb
R^2 = 0.80
Read Length Group = >=4000 kb
R^2 = 0.80
Read Length Group = >=5000 kb
R^2 = 0.74
Read Length Group = >=6000 kb
R^2 = 0.68
Read Length Group = >=7000 kb
R^2 = 0.60
Read Length Group = >=8000 kb
R^2 = 0.52
Read Length Group = >=9000 kb
R^2 = 0.33
Read Length Group = >=10000 kb
R^2 = 0.07
telomere_length
read_length
d
Read Length Group = >=0 kb
R^2 = 0.28
Read Length Group = >=1000 kb
R^2 = 0.28
Read Length Group = >=2000 kb
R^2 = 0.27
Read Length Group = >=3000 kb
R^2 = 0.26
Read Length Group = >=4000 kb
R^2 = 0.23
Read Length Group = >=5000 kb
R^2 = 0.21
Read Length Group = >=6000 kb
R^2 = 0.19
Read Length Group = >=7000 kb
R^2 = 0.15
Read Length Group = >=8000 kb
R^2 = 0.10
Read Length Group = >=9000 kb
R^2 = 0.08
Read Length Group = >=10000 kb
R^2 = 0.08
telomere_length
read_length
e
Density
Read Length (bp)
Long-Read Sequencing Modality
ONT
PacBio
f
Subtelomere Length (bp)
EcoRV
HinfI/RsaI/EcoRV
Long-Read Sequencing Modality
ONT
PacBio

**Fig. S4. PacBio but not Oxford nanopore telomere length measurements are bottle-necked by read length.** HEK293T telomere length distributions representing sample of 1000 telomere measurements randomly selected from telomeric reads with a read-length longer than incrementally increasing cutoffs, as measured by PacBio *(*a, *26)* or ONT long-reads (b, original data). Correlation between measured telomere length and read length from PacBio (c, *26*) or ONT (d, original data) long-read sequencing digital telomere length measurement. (e) Read length density distributions for PacBio or ONT data previously demonstrated. (f) Aggregate subtelomere length measured following single or combination restriction digestion in telomeric reads by ONT (left, original data), or PacBio (right, *26*).

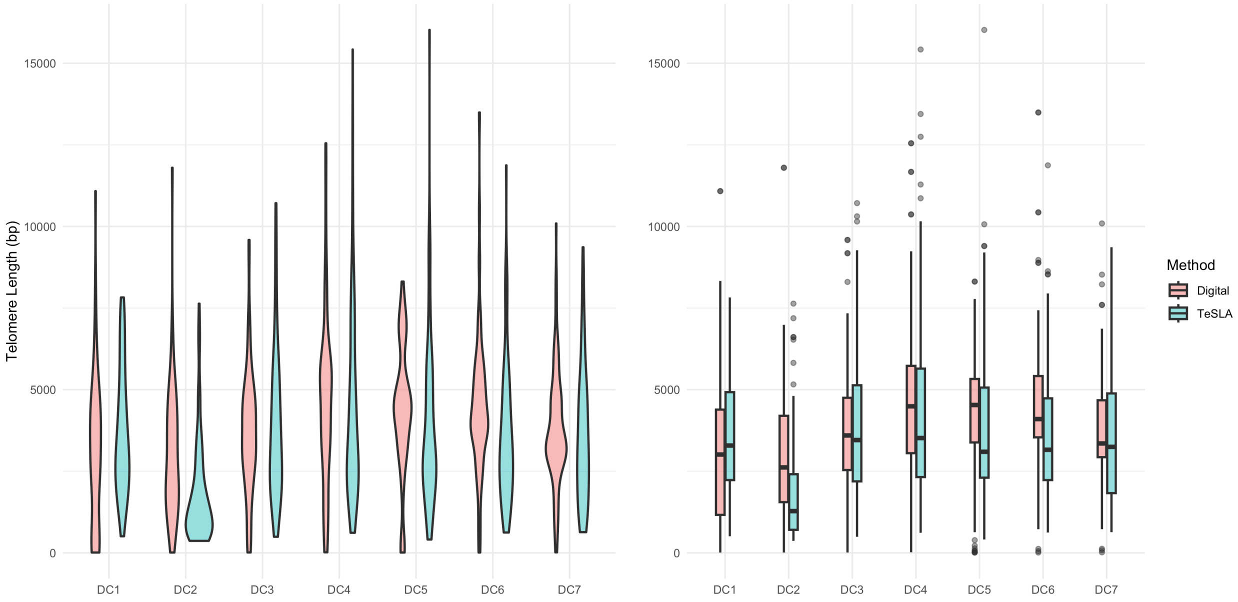


**Fig. S5. Comparison of digital telomere measurements and in-gel measurements of TeSLA Southern blot bands from RTEL1 variant cohort PBLs.** Violin (left) and boxplot (right) of digital telomere measurements by long-read sequencing (red) and in-gel measurements of TeSLA

Southern blot bands (blue) from RTEL1 variant cohort PBLs. TeSLA measurements originally published in (*22*).

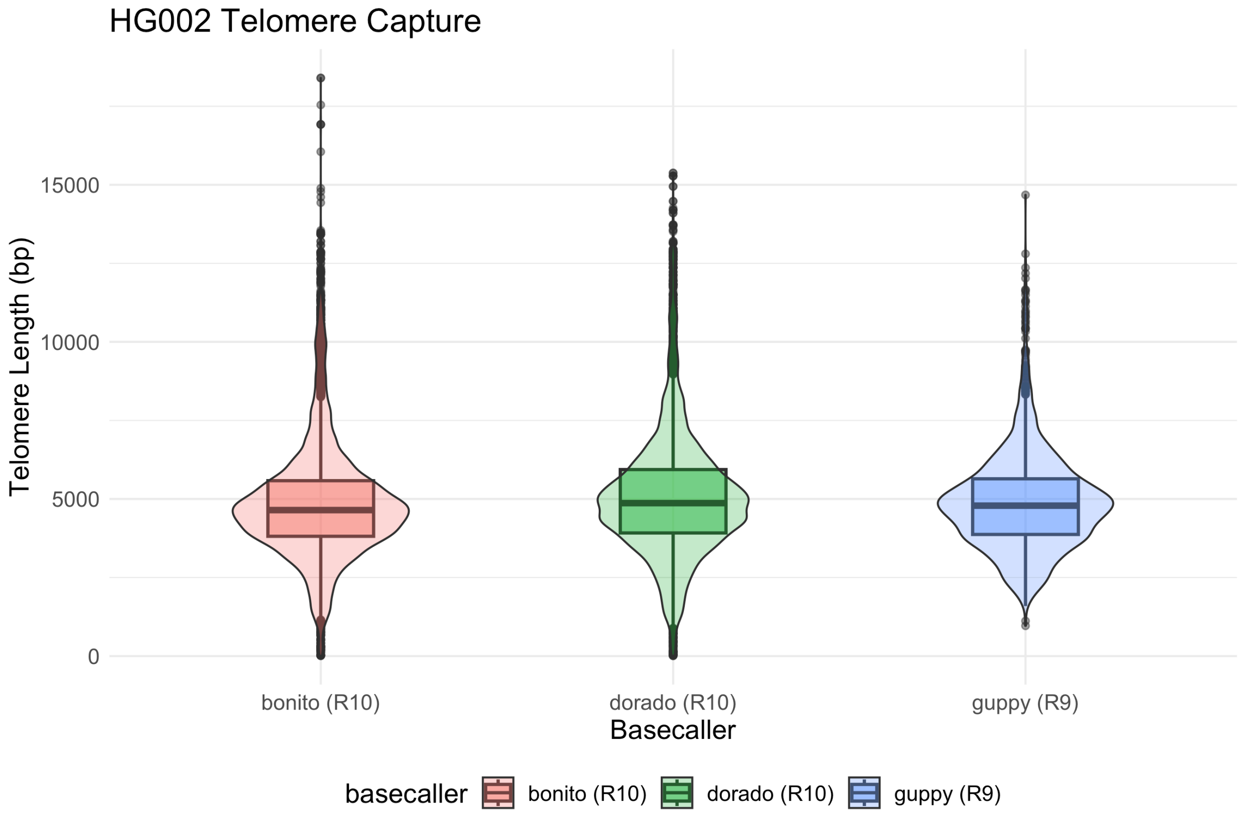


**Fig S6. HG002 digital telomere measurement and basecaller performance comparison.** Telomere capture sequencing and digital measurement across two technical replicates (R9 or R10 sequencing chemistry) and three basecalling models. Bonito (custom ONT model HG002.k1, n=10465) and dorado (v0.3.4, dna_r10.4.1_e8.2_400bps_sup@v4.2.0, n=10353,) models were used to basecall the same raw sequencing data. Guppy (v6.5.3, n=2122) R9 basecalls represent a technical replicate. Telomere measurement was performed with Telometer in all cases.

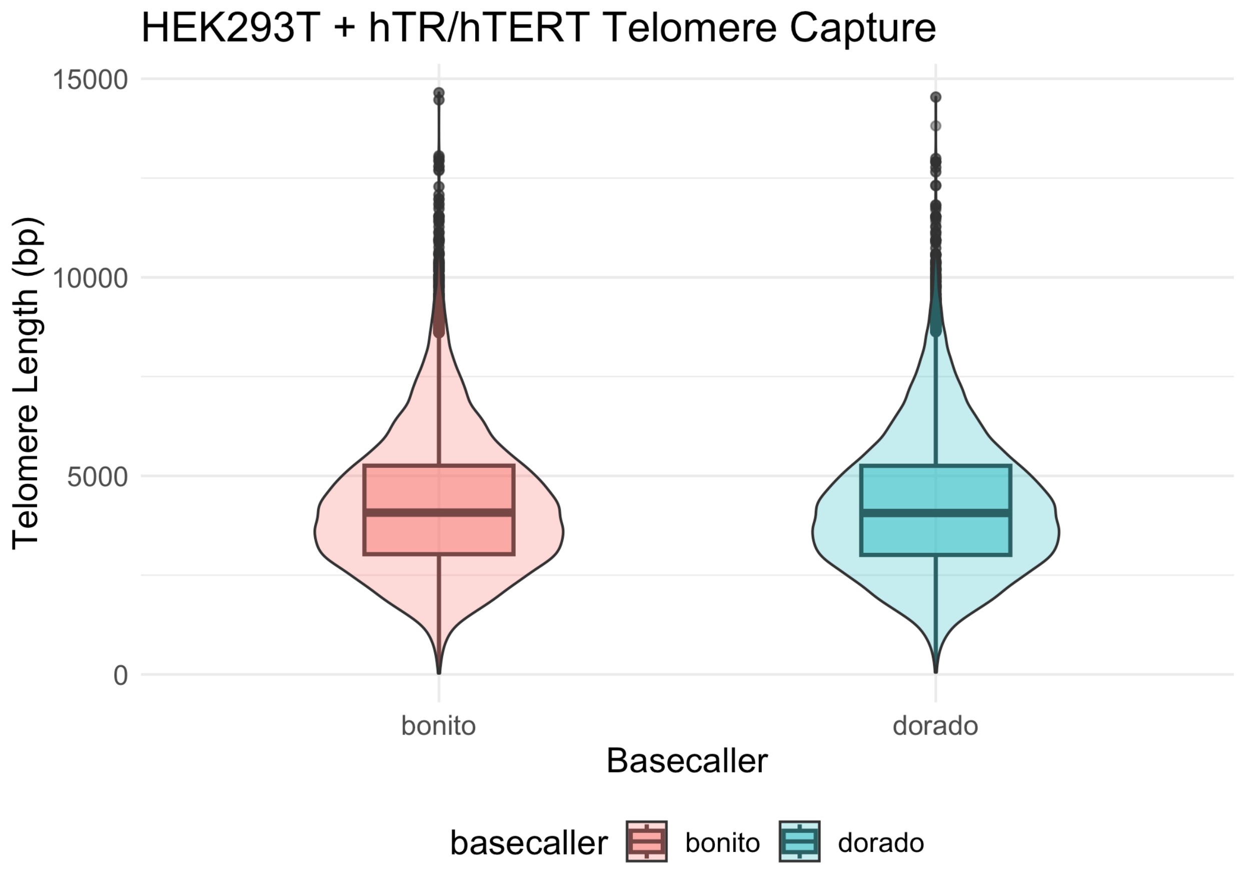


**Fig S7. HEK 293T digital telomere measurement and basecaller performance comparison.** Telomere capture sequencing and digital measurement across two basecalling models. Bonito (custom ONT model HG002.k1, n=21402) and dorado (v0.3.4, dna_r10.4.1_e8.2_400bps_sup@v4.2.0, n=21282,) models were used to basecall the same raw sequencing data. Telomere measurement was performed with Telometer in both cases.

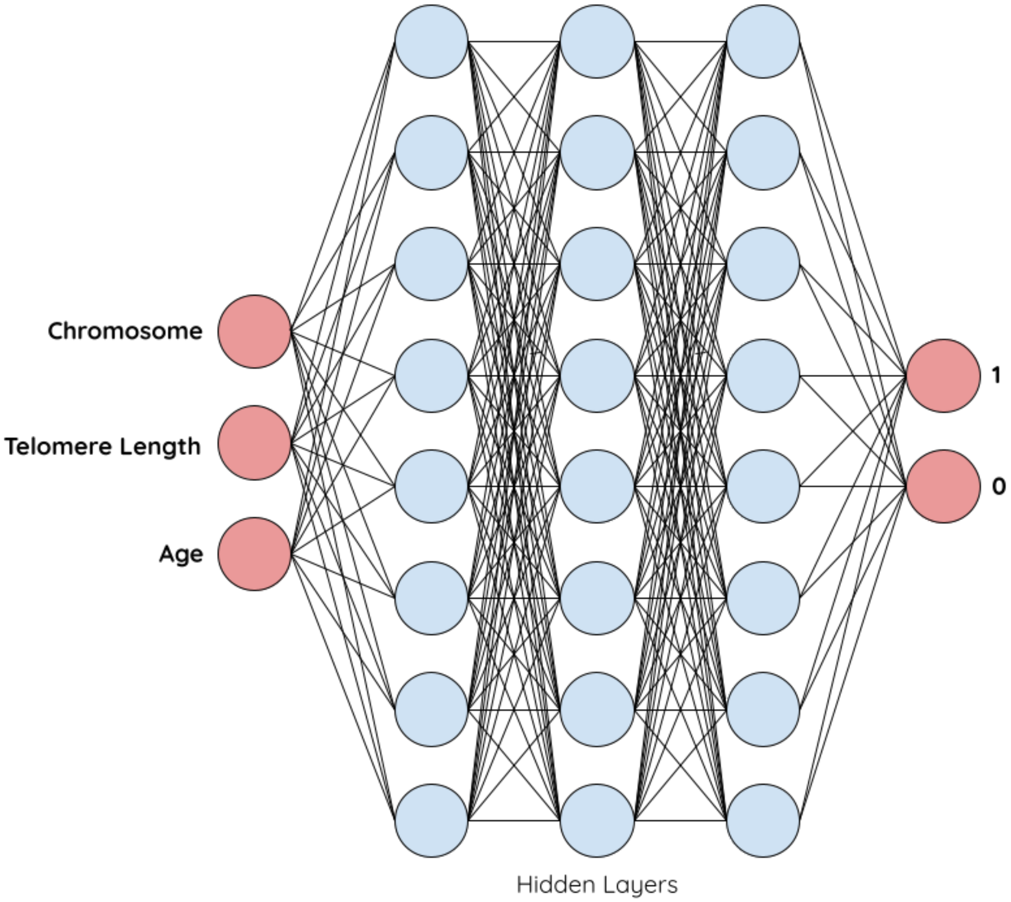


**Figure S8. MLP binary classifier model architecture.** Telometer output is used as input and three separate models were trained for three binary classification outputs: healthy versus diseased, healthy versus asymptomatic carrier, and healthy versus diseased or asymptomatic carrier.
